# Network controllability measures of subnetworks: implications for neurosciences

**DOI:** 10.1101/2022.09.11.507468

**Authors:** Julia Elina Stocker, Erfan Nozari, Marieke van Vugt, Andreas Jansen, Hamidreza Jamalabadi

## Abstract

Recent progress in network sciences has made it possible to apply key findings from control theory to the study of networks. Referred to as network control theory, this framework describes how the interactions between interconnected system elements and external energy sources, potentially constrained by different optimality criteria, result in complex network behavior. A typical example is the quantification of the functional role certain brain regions or symptoms play in shaping the temporal dynamics of brain activity or the clinical course of a disease, a property that is quantified in terms of the so-called controllability metrics. Critically though, contrary to the engineering context in which control theory was originally developed, a mathematical understanding of the network nodes and connections in neurosciences cannot be assumed. For instance, in the case of psychological systems such as those studied to understand the psychiatric disorders, a potentially large set variables are unknown. As such, while the measures offered by network control theory would be mathematically correct, in that they can be calculated with high precision, they could have little translational values with respect to their putative role suggested by controllability metrics. It is therefore critical to understand if and how the controllability metrics computer over subnetworks would deviate, if access to the complete set of variables, as in neurosciences, cannot be taken for granted. In this paper, we use a host of simulations based on synthetic as well as structural MRI data to study the potential deviation of controllability metrics in sub-compared to the full networks. Specifically, we estimate average- and modal-controllability, two of the most widely used controllability measures in neurosciences, in a large number of settings where we systematically vary network type, network size, and edge density. We find out, across all network types we test, that average and modal controllability are systematically, either over- or underestimated depending on the number of nodes in the sub- and full network and the edge density. Finally, we provide a formal theoretical proof that our observations generalize to any network type and discuss the ramifications of this systematic bias and potential solutions to alleviate the problem.

## 1 Introduction

Characterized by a set of elements and their connections, networks are ubiquitous in neurosciences ^1^. On multiple levels of abstraction, the brain can be studied as a complex system where the interaction of a large number of elements, across many scales, enables the emergent properties referred to, collectively, as brain dynamics ^2^. Structural networks based on structural connectivity ^3^, functional networks based on functional and effective connectivity ^4^ and structural covariance based on the interrelation of gray matter properties ^5^ are among the most studied networks in neurosciences. Similarly, networks are being increasingly used to study cognitive and psychological constructs ^6^. In fact, network theory of psychopathology posits that mental disorders can be best conceptualized and studied as causal systems of mutually reinforcing symptoms ^7^.

Importantly, most networks studied in neurosciences concern fundamentally dynamic processes that change over time. For instance, a network model of a mental disorder is most relevant to inform the clinician if it can capture how the symptoms temporally unfold and how they functionally affect each other. While a large number of statistics is developed to study networks in general (see Gosak et al.^8^ for a review), this and similar questions are being increasingly approached using results from network control theory ^9,10^. With close ties to the dynamical systems theory, this framework concerns how the interactions between interconnected elements, potentially constrained by a set of given optimality criteria, result in complex systemic behavior ^11^. Central to network control theory is the idea that in a system of interacting components, the change in one element causes a cascade of events in the whole system and that these changes follow a general mathematical structure ^11–13^. When applied to neuroimaging data, network control theory provides a novel mechanistic framework to describe neural activity and the nodes that mostly drive the temporal dynamics^14,15^. Such a dynamical approach lets us quantify the disposition of nodes to change the system rather than gaining a momentary snapshot from centrality metrices. A typical example is quantification of the role that each brain region or symptom plays to shape the temporal dynamics of the brain activity or the clinical course of a disease ^16^. For this and similar questions, assuming linear temporal dynamics, network control theory provides a set of measures that mathematically quantify the relevance of the nodes ^17^. Specifically, average-, and modal-controllability are among the most widely measures used in neurosciences to study neural ^9,10,18–20^ as well as psychological symptom dynamics^21,22^ and are suggested to measure the overall power of nodes to affect network dynamics (see Karrer et. al. for a detailed mathematical introduction ^23^).

Despite the initial success and increasing popularity of employing network control theory in neurosciences, the statistical properties and reliability of these measures have remained poorly understood. Importantly, contrary to the engineering systems for which control theory was originally developed, a complete and agreed-upon understanding of the networks, nodes, and connections that are most relevant for each problem in neuroscience is far from available. For instance, in the case of psychological networks where nodes can resemble psychological states and their interactions, even the variables that affect the phenomenon under study are only partly known ^24^. In fact, since most of the psychological variables are interrelated (see Robinaugh et al.^25^ for a current review), no network can be assumed to be complete, as more nodes can always be added to any network^25^. Similarly, in many studies involving neuroimaging data, the choice of the parcellation atlas, inclusion of subcortical regions in the analysis, the resolution of analysis (e.g. voxel based vs regional analysis), and even the regions to be included in the analyses (the so-called region of interest analysis ^26^) follow heuristics and statistical considerations rather than fundamental derivations. It is therefore critical to understand if, and how, the “estimated” controllability metrics that are computed from networks consisting of only a subset of all possible nodes deviate from their nominal values should they were computed over a complete network (see Figure 1 for an illustrative example). To address this issue, we use a host of simulations based on synthetic data (see Figure 1 for an example) as well as data from brain structural connectome to study the statistical properties of controllability metrics in subnetworks. We replicate all simulations in three canonical network models, producing networks with different sizes, edge density, and number of nodes to compare the controllability metrics in the full- and in the sub-networks and finally provide a mathematical formulation for our main finding. -

**Figure 1:**
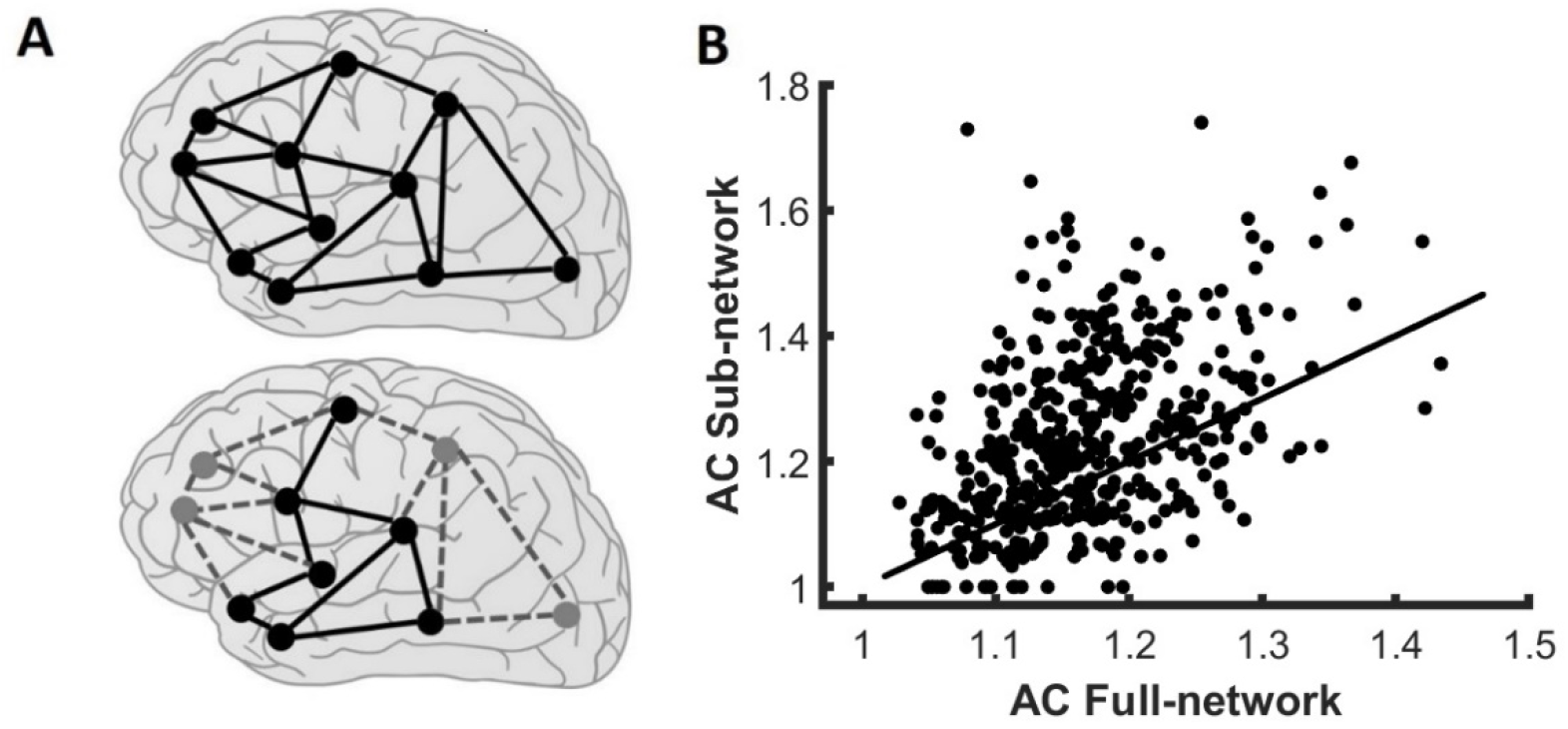
Controllability metrics are biased if only part of the network is included in the analysis. A) a representing example of a full network (above) and one subnetwork (below). Although not included in the calculations, the connection to other regions would still affect the controllability metrics of the nodes in the subnetwork. In this manuscript, we study ifthe deviation from the nominal values (i.e., those computed from the full network) systematically bias the estimations or shows itself as a zero-mean noise. (B) A simulated example of the deviation in controllability metrics. In this example, we simulated random graphs with 50 nodes with low connection density (see Methods) and compare the average controllability of the nodes with the estimated value when only a subnetwork with one fifth of the nodes is selected. The solid line represents the identity line. In the example here, we notice that the average controllability is mostly overestimated (the values lie mostly above the identity line), and the deviation seems to be largest for the nodes with highest original average controllability.

## 2 Methods

### 2.1 Network generation and subnetwork sampling

A network, also called a graph, is mathematically defined as *G* = (*V, E*) where *V* = {*V*_1_,…, *V*_*N*_} represents the set of nodes (e.g. brain regions, set of symptoms, etc.) and *E* = {(*V*_*i*_,*V*_*j*_)|(*i,j*) ∈1:*N*} represents the set of edges connecting the nodes (e.g. white matter tracts, causal links across symptoms, etc.). The connections could be binary or weighted (different connections are similar or have different strength) and could be either directed or undirected (the connection between different nodes could have different strength in each direction, for instance, node *V*_1_ can influence *V*_2_ but not vice versa). A graph is therefore an abstraction of the relation among the constructing components of a system and is often used to relate the function to the structure of a system. Although every real graph is different from any other, it is useful to study the graph properties in families of graphs that have similar statistical properties. Following previous work on the assessment of graph properties, we simulate three canonical network types that are shown to be most representative of psychological and neural phenomena ^27,28^.

1. **Random networks**, also called Erdős–Rényi (ER), are characterized by a fixed independent edge probability. That is, for a graph with *N* nodes and an edge probability of *p*, each edge exists with a probability of *p* and as such, the nodal degrees (the number of connections for each node) follow a binomial distribution with the expected value of 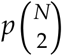. Due to its property of being randomly interconnected, the random graph can be easily used as a baseline to compare to other network types. In this paper, we generate the networks using the codes available from the MATLAB Central File Exchange ^29^.
2. **Small world networks** are characterized by the small distance between any two randomly chosen nodes in the graph and exhibits low short path length and low clustering coefficient. In this paper, we use the generation algorithm proposed by and named after Watts-Strogatz. For a graph with *N* nodes, this model has two parameters of *K* and *β* and that specify the mean degree and rewiring probability, accordingly. Small world networks result in highly interconnected regions of nodes, called hubs that are sparsely interconnected. We generate the networks using built-in MATLAB function WattsStrogatz.
3. **Scale-free networks** are characterized by the exponential distribution of the degree. In this paper, we use the scale-free network proposed by and named after Barabasi that has two parameters of *m*_*o*_ and *m* that specify the seed and average degree. Scale-free models have a less hub-like structure than a small-world and are suggested to most closely resemble biological systems ^30^. The implementation of the models is done based on the codes available from the MATLAB Central File Exchange^31^.

In our simulations, for each network type, we generate undirected binary graphs. This choice is motivated by 1) the fact that binary networks are ubiquitous in neurosciences and have been shown to form a valuable model of the reality ^32,33^, 2) the controllability metrics behave similarly to weighted networks ^28^, 3) simulation of binary networks requires less number of parameters and the interpretation is clearer. However, to make sure that our results generalize to realistic settings, we compare the results based on synthetic binary networks to the brain structural data modeled by weighted network (section 3.3) and further provide formal theoretical proof for our main observations that is independent of the network types (section 3.4). In our simulations, we vary the number of nodes *N* between 10 and 200 and the mean degree between 0.1 × *N* to 0.1 × *N*. By changing the mean degree, we aimed to include the full spectrum of sparse and densely connected networks as they can have fundamentally district statical properties^8^. For each full network, we subsampled nine subnetworks from which we selected a subset of the nodes (starting from one fifth, minimally 2 nodes for a full network of size 10) and their associated edges by following an iterative procedure: in the first step, we randomly selected one fifth of the nodes using the randperm function, integrated in MATLAB R2021b. For each new subnetwork with more nodes, we similarly selected random additional nodes from the whole set of nodes. Thereby each subnetwork builds on the previous node selection. In doing so, we made sure that the subnets with varying size are comparable in that they include incrementally bigger networks that encompass the previous smaller subnetworks. We repeated the whole procedure 100 times to account for randomness in the selection process. All simulations were done in MATLAB 2021b.

### 2.2 Real Dataset and comparable simulation data

To compare our simulations to real-world data, we use brain imaging data originally acquired and described by Muldoon and colleagues that is publicly available and preprocessed according to the best practices in the field ^34^. In short, eight subjects (mean age 27 ± 5 years) were scanned three times. In each session, they undergo Diffusion Spectrum Imaging (DSI), along with T1 anatomical scans. The DSI scans are sampled in 257 directions and are reconstructed in DSI studio using *q*-space diffeomorphic reconstruction (QSDR). More details on the data preprocessing can be found elsewhere ^34^. Structural brain networks are then obtained by subdividing the entire brain into 83 anatomically distinct brain areas (network nodes). For each of these networks with 83 nodes, we then randomly select subnetworks (20-100%), similar to our simulations. This time we use the randsample function to choose appropriate node indices of the given networks. Then we iteratively select additional nodes for each of the nine subnetworks. This subnetwork selection is repeated for all subnetwork sizes 20 times. In order to make simulation data comparable to the real dataset, we additionally simulate networks of a similar network size (N=83) and mean degree. We choose the mean degrees of those simulations according to the average degrees of all subsets of the real data.

### 2.3 Network Controllability Metrics

Following the mainstream literature in studying controllability properties of brain networks ^9^, we assume a noise-free discrete-time linear time-invariant (LTI) dynamical system where the interaction terms (i.e. networks) are built as described above. Within the framework of network control, controllability measures refer to the importance of single nodes in steering the network’s dynamics. In particular, average and modal controllability have been suggested to quantify the nodal ability to execute easy and difficult state transitions ^23^, respectively, and have been shown to be sensitive enough to relate to a wide range of brain cognitive and functional properties ^16^. Mathematically, average controllability (AC) of node *k* is defined as:

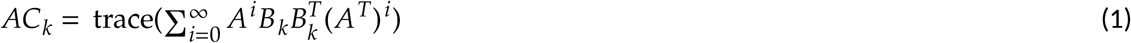

where *A* is the network under study and *B* the *k*^*th*^ canonical vector (a vector of length *N* with all zero elements except a 1 in its *k*’th position). Modal controllability (MC) is calculated by:

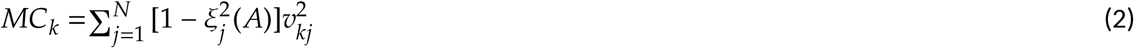

where *ξ*_*j*_ is the *j*’th eigenvalue and *v*_*kj*_ is the *k*’th element of the *j*’th eigenvector of *A*. Since average controllability is always above 1 and modal controllability below 1, for each original network *A* and each subnetwork *A*_*s*_, we define the deviations in *AC*_*s,k*_ and *MC*_*s, k*_ (computed over the subnetwork) as follows:

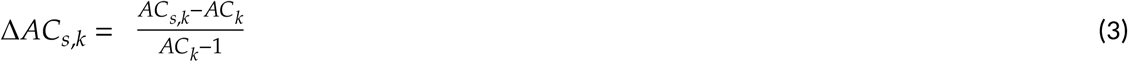

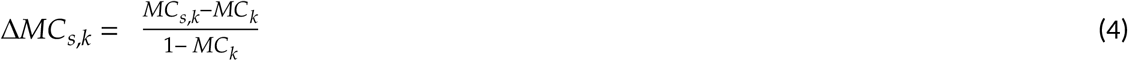

## 3 Results

### 3.1 Deviation of estimated controllability metrics depends on nodal degree

Controllability metrics are closely related to nodal degree ^15^. As such, we begin by studying the deviation of estimated controllability metrics in nodes with different degrees. Following the procedure outlined in Methods, we simulated examples of three network families and vary network sizes while keeping the mean degree and the ratio of nodes in the subnetworks constant. For each node and each graph, we calculate the difference in controllability metrics between the full and sub-networks (see equations 3-4). Our results show that, similarly across all network types, modal controllability is systematically underestimated, and the underestimation is more prominent for the nodes with higher degree. Average controllability on the other hand shows a more complex behavior. It is on average underestimated for nodes with low degree, but it is overestimated for nodes with higher degree. Both effects tend to become larger for the smaller networks and for the nodes with higher degree (Figure 2). Interesting though, our results also show that average controllability is generally less biased compared to modal controllability for a wide range of network parameters.

**Figure 2:**
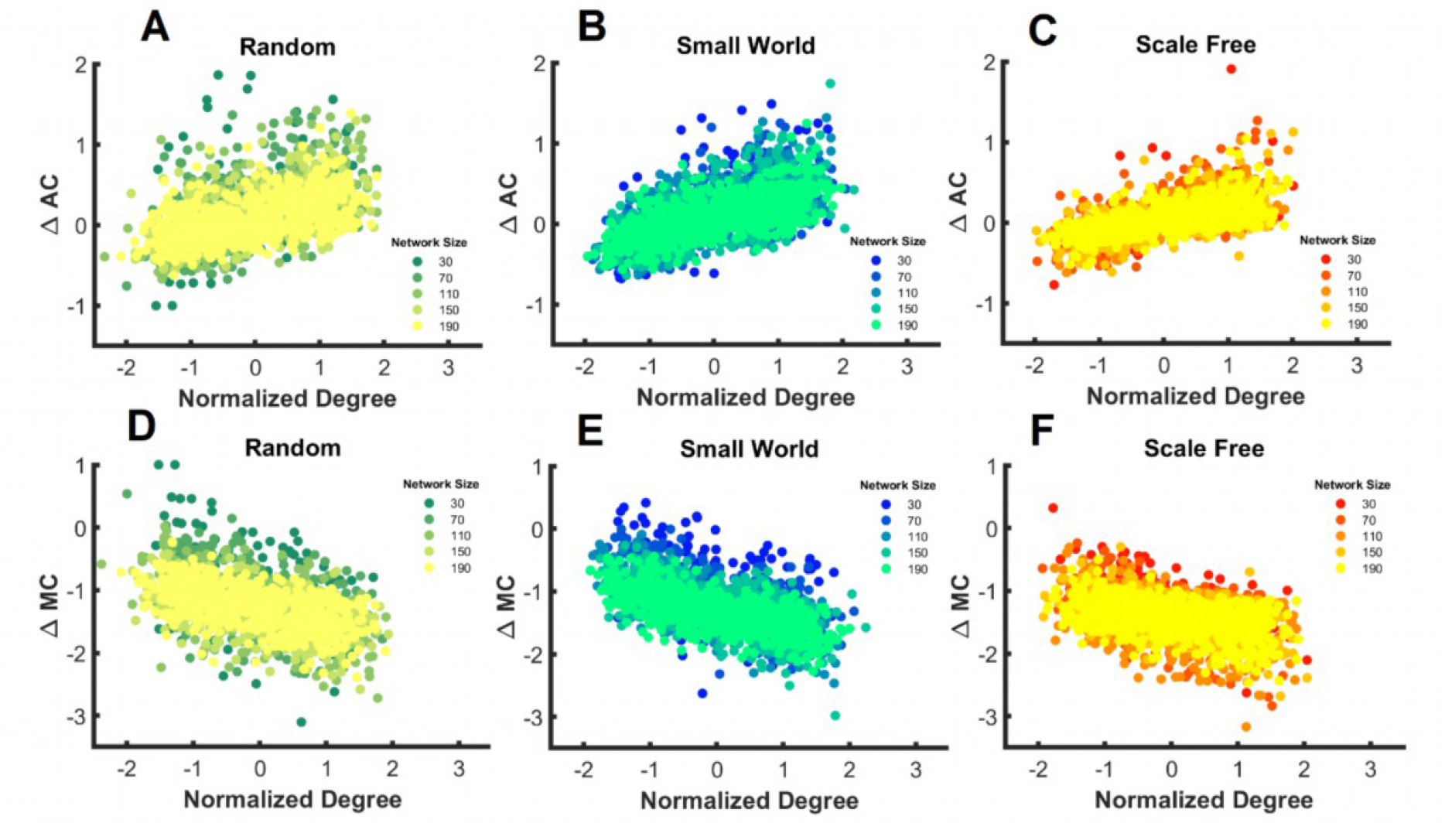
Deviations of controllability metrics relate to the normalized nodal degree (z-scored nodal degree). We simulated networks with varying sized while keeping the mean degree (= N × 0.57) and the ratio of nodes in the subnetworks constant (N × 0.4 see Methods for details). A-C) AC is underestimated for nodes with small and overestimated for nodes with high degree. D-F MC is underestimated for all nodes, but the bias is most prominent for the nodes with highest degree.

Having established that the controllability metrics systematically deviate in subnetworks, we further studied how this deviation relates to the edge density. For this, we simulated networks with varying mean degree (see Methods for details). Our results show that the modal controllability is systematically underestimated across all network types, and the deviation is comparable across all values of edge density. Average controllability is however overestimated for graphs with low edge density and underestimated for graphs with high mean degree (see Figure 3). Interestingly, similar to the results in the previous section, we observe that across a large number of parameters, average controllability is less biased than modal controllability.

**Figure 3:**
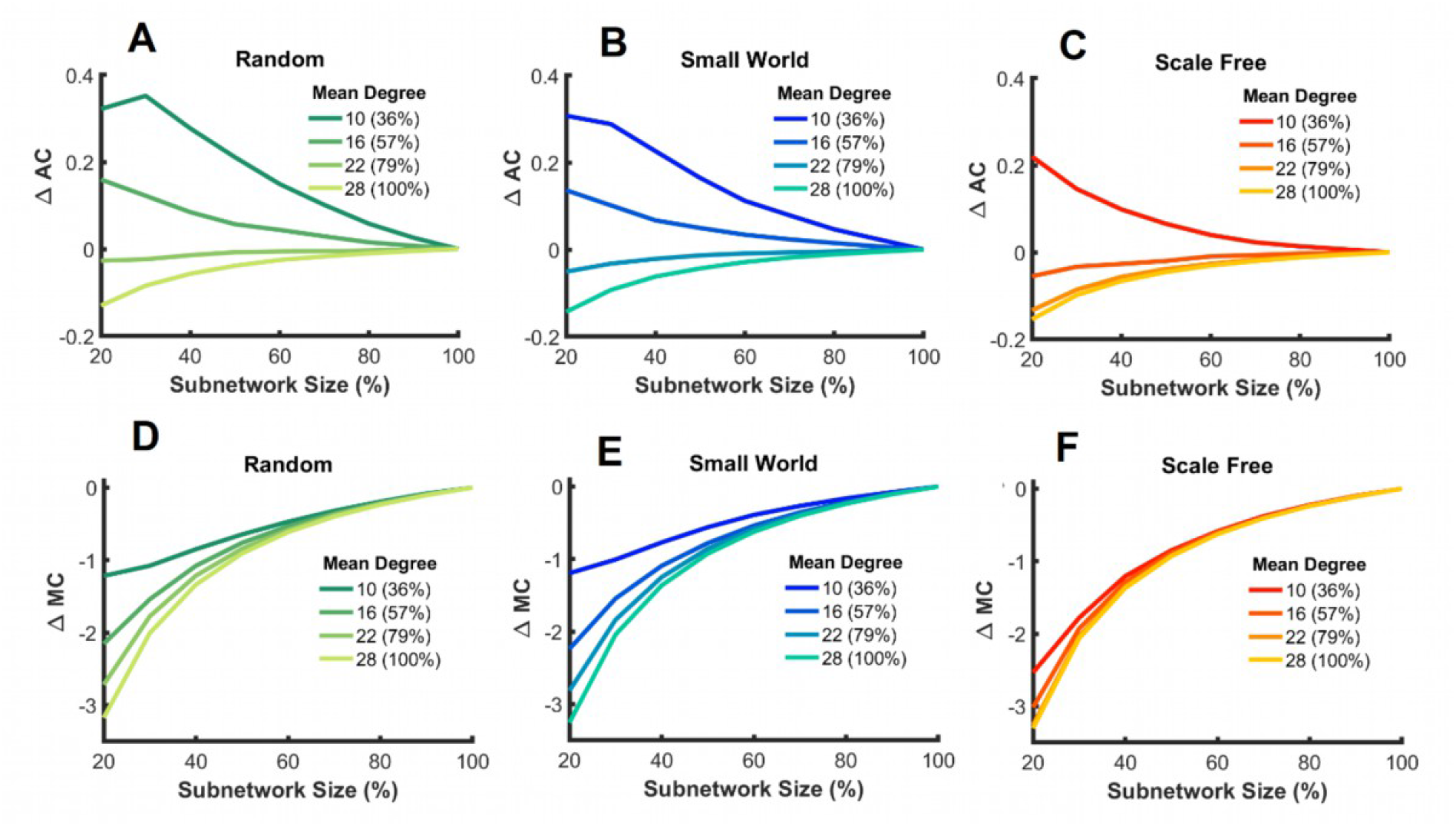
Deviation of controllability metrics for networks with varying mean degree. A-C) Deviation of average controllability across. D-F) Modal controllability is independent of the edge density underestimated.

### 3.2 Deviation of estimated controllability metrics depends on network size

Previous results demonstrate that the controllability deviation depends on the size of the selected subnetwork. We further asked, if beyond the selected number of nodes (subnetwork size), the size of the full network would be an additional independent factor that affects the controllability deviation. To address this question, we simulate networks with a fixed mean degree and estimate the controllability deviation while systematically changing the size of the networks and subnetworks. Our results show that average controllability is systematically over, and modal controllability is systematically underestimated (see Figure 4), but the deviation is largest for smaller subnetworks. The size of the full network seems to be less relevant for the controllability deviation and is different between the controllability metrics where larger network sizes are associated with higher deviation of average and lower deviation of modal controllability. In line with our previous simulation, we also note that, the deviation of modal controllability is much larger than average controllability (up to five times).

**Figure 4:**
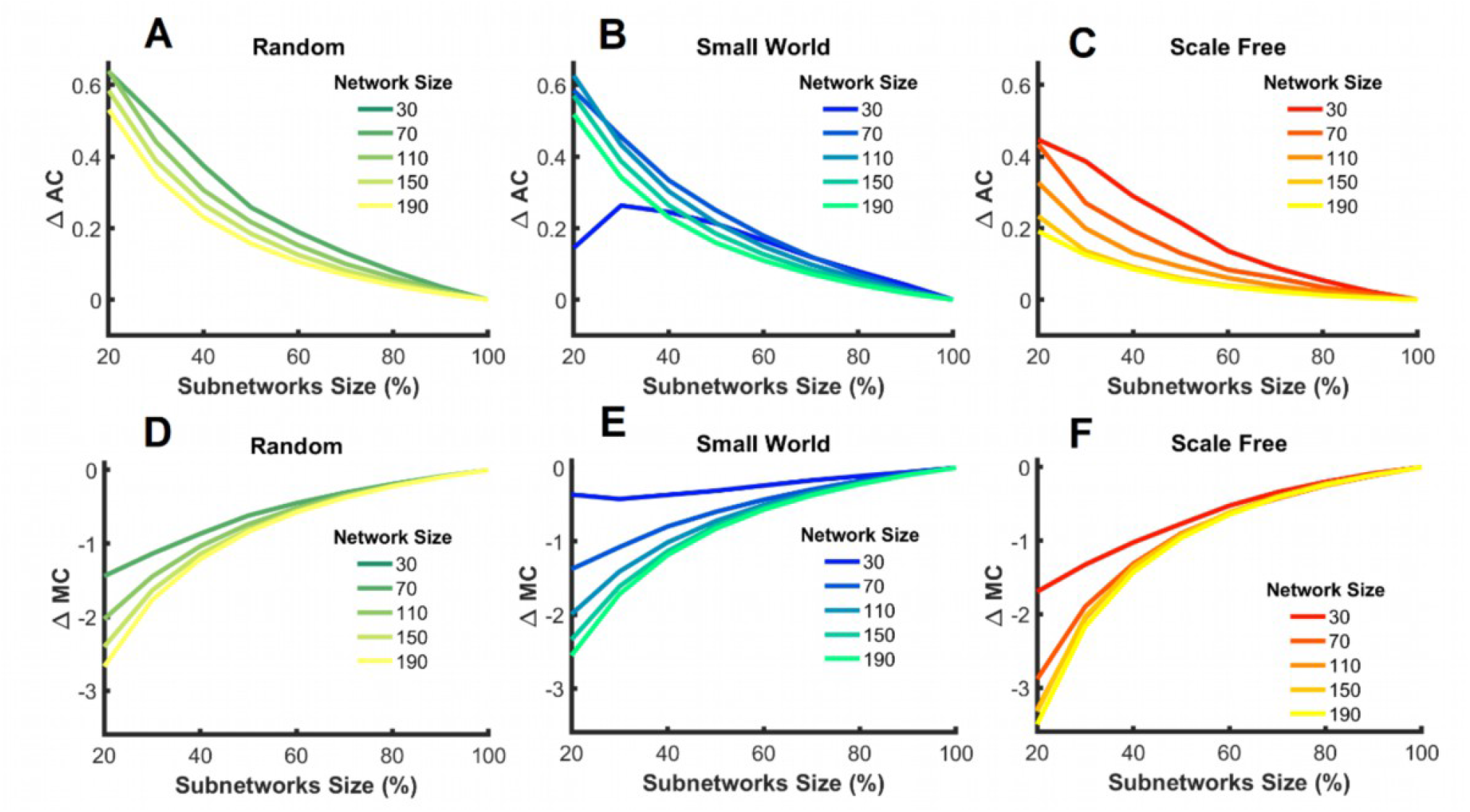
Controllability deviation for different network sizes. A) Average controllability is systematically overestimated and B) Modal controllability is systematically underestimated across all network types. The deviation is highest for small networks and for modal controllability. It is interesting to note that for small world networks, one sees that the deviation for average controllability for very small number of nodes does not follow the same pattern as the other networks. This, we believe, relates to the instability of the network generation for very small sizes.

### 3.3 Deviation of estimated controllability metrics in brain networks

Finally, we tested if the deviation we estimate from our simulations aligns with real data. To do this, we used data from brain structural connectome of 8 subjects in 3 sessions each with 83 brain regions (see Methods). We simulated networks comparable to our simulations, having the same number of nodes and mean degree and from this, we sampled 9 different subnetworks. The results show that the deviation in average and modal controllability is similar to our models.

### 3.4 Theoretical results

In previous sections, we experimentally established, using synthetic as well as neuroimaging data, that average controllability is mostly over- and modal controllability is mostly understimated (Figures 2-5). Here, we provide a formal theoretical proof that this result holds for any network. Consider, for simplicity, random symmetric matrices *A*_*N*×*N*_ with independent (modulo symmetry) and identically distributed edge weights. By the well-known semi-circle law for the spectra of random matrices ^35^ if one keeps the distribution of edge weights independent of the network size, the largest eigenvalue of *A* grows as *N* while the remaining *N* − 1 eigenvalues of *A* (a.k.a. the eigenvalue “bulk”) grow as √*N*. Therefore, once we normalize A by dividing it by its largest eigenvalue (as almost always done to ensure stability in controllability analyses ^9,16^), the bulk of eigenvalues of *A* decays as 1/√*N*. Therefore, for large networks (dozens of nodes or more), normalized *A* has one eigenvalue equal to 1 and the rest of eigenvalues near 0. Now, by the Cauchy’s Interlace Theorem for eigenvalues ^36^, the eigenvalues of *A*_*s*_ “interlace” (lie between pairs of) the eigenvalues of *A*. Importantly, this means that the largest eigenvalue of *A*_*s*_ (call it μ_1_) lies between 1 and the bulk of eigenvalues of *A*, while the remaining (bulk) of eigenvalues of *A*_*s*_ lie between the bulk of eigenvalues of *A*. Since the bulk of eigenvalues of *A* are small (order 1/√*N*) and all lumped together, so are the bulk of eigenvalues of *A*_*s*_. Therefore, on average, the bulks of *A* and *A*_*s*_ are nearly indistinguishable. However, since *A*_*s*_ is re-normalized to make its largest eigenvalue equal to 1, all of its eigenvalues are multiplied by 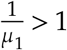, hence boosting its bulk above that of *A*. Thus, after normalization, both *A* and *A*_*s*_ have one eigenvalue equal to 1 and one bulk of small eigenvalues, where the bulk of *A*_*s*_ is an order of 1/*μ*_1_ larger than that of *A*. This fact nicely explains the mainly negative (positive) trends we observed in Δ*MC*(Δ*Ac*) since, as we showed in our recent work ^18^, the average (over all network nodes) AC and MC have the simple expressions as in equations 5-6.

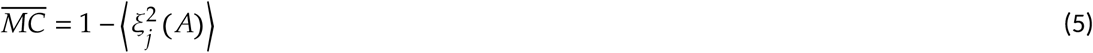

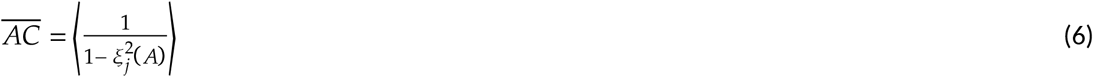

where ⟨.⟩ denotes average over all *j*. In words, average MC is negatively correlated with mean squared network eigenvalues (and hence smaller in subnetworks), while average AC is positively correlated with mean squared eigenvalues (and hence larger in subnetworks).

## 4. Discussion

In neuroscience, network control theory is increasingly used, for good reasons ^10,18,19,21,37^, to study neural, biological, and psychological constructs. It relates fundamental theory-driven results from controls literature to the study of networks that are the natural points of interest in neurosciences^14^. However, as with other computational approaches, the added value of such synergies can only be fully exploited once the statistical properties and theoretical prerequisites have been thoroughly investigated ^15,38,39^. Here, building upon the notion that biological and psychological networks are fundamentally different from the systems studied in the engineering context, we studied if the estimated controllability metrics would manifest any systematic deviation from their nominal value. Simulating a large number of networks of varying types and sizes, we found that average and modal controllability, two of the most widely adopted controllability metrics in neuroscience^23^, are systematically biased and that from these two, average controllability seems to be the one with lower bias. Indeed, as Figure 5B illustrates, in a setting similar to those often used in neuroimaging studies, the deviation in average controllability is up to three times smaller than that of modal controllability. Importantly, we further provided theoretical results that formally supports our empirical observations independent of network model, size, and edge density.

**Figure 5:**
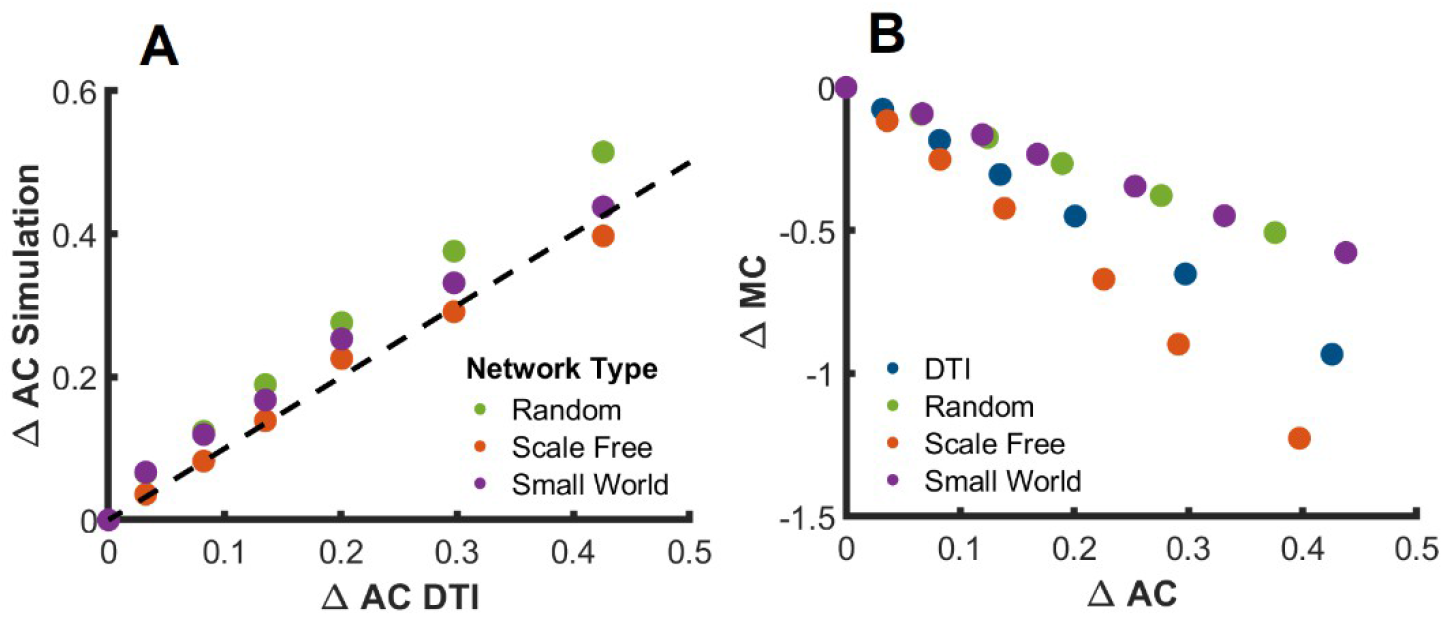
Comparing deviations of controllability measures of brain networks to simulations. (A) Deviation of average controllability for the brain structural connectome compared to the simulations. Here we built networks with 83 nodes (as in the brain structural connectome, see Methods for details) with the same mean degree as in the brain data. We sampled nine subnetworks where we varied the subnetwork size between 20% to 100% of the original nodes. The dashed line shows the identity line. (B) illustrates deviation of average and modal controllability for different subnetworks (20%-100%) from whole networks with N=83. The average deviations of all networks (Random, Scale Free and Small World, as well as DSI) are depicted together with the interindividual averages across scans.

Furthermore, beyond the network size, nodal degree seems to be an important factor in the size of the deviation. Since nodal degree is also independently related to the controllability metrics^9,15^, our results suggests that any statistical test comparing controllability metrics should correct for the effect of nodal degree. Otherwise, the associated effects could be a result of inflated controllability metrics or the degree itself rather than the topological properties embodied by controllability metrics. Relatedly, our results suggest the controllability metrics for larger subnetworks are less deviated. While this is mathematically a trivial phenomenon, it has important ramifications in practice. In the context of neuroimaging studies, whole-brain analysis is more likely to be accurate and in the context of psychological networks, including more items is likely to benefit the robustness of the metrics. Yet, larger networks also require more data to reliably estimate (e.g., Epskamp et al.^40^), pointing to a trade-off that every researcher will need to consider in their experiment design. Related, our observations in this paper do not directly undermine the application of controllability metrics when the same metric is compared across the nodes within the same network. However, since no two networks are completely identical, and different controllability metrics are associated with different biases, our results warrant caution when tests are carried out across networks or metrics.

Finally, a note is warranted on system identification and the various ways in which network models can be extracted from data. A common practice in the applications of network control theory in neuroscience is to obtain the adjacency matrix *A* from structural connectome data extracted from diffusion imaging or based on temporal correlations, which although attractive from many perspectives, the models are then arbitrarily weighted. As shown in the formal derivation above (section 3.4), the common normalization of the *A* matrix plays a major role in the bias we empirically observe in subnetwork-estimated controllability metrics. This normalization, importantly, is neither necessary nor recommended from a pure controls perspective – all controllability metrics can be computed for stable systems (even if the largest eigenvalues is not exactly one) and many (using finite-horizon controllability Gramians) are computable for unstable systems. However, the arbitrary weighting of the networks makes normalization almost inevitable. Networks learned using the system identification theory ^41^, on the other hand, have well-defined edge weights computed from nodel time-series data regardless of the selection of network nodes from which data has been recorded. This can in turn significantly improve the reliability of controllability metrics and limit the effect of unmeasured nodes to an increase in the unexplainable variance of model predictions.

## 4 Conclusion

Our results warrant caution when using engineering motivated metrics in neurosciences. Network control theory has a lot to offer but the many differences between engineering systems for which this framework is developed and neurosciences must be properly addressed. Further, it is worth highlighting that we here analyzed average and modal controllability only due to their prevalence in the literature, but network controllability provides a significantly broader picture than these metrics. Future work should analyze similar (sub)network size effects on other controllability metrics.

## Availability of data

The codes to replicate the simulations will become available publicly upon acceptance of this paper. The DSI data used in this study is publicly available at https://datadryad.org/stash/dataset/doi:10.5061/dryad.8g4vp.

## Author contribution statement

Conceptualization: ES, HJ, AJ. Methodology and Validation: HJ, ES, EN. Simulations and analysis: HJ, ES, EN. Writing: all.

